# MiRNA expression profiles of serum exosomes derived from individuals with latent and active tuberculosis

**DOI:** 10.1101/316794

**Authors:** Lingna Lyu, Jinghui Wang, Hongyan Jia, Liping Pan, Zihui Li, Fengjiao Du, Boping Du, Qi Sun, Zongde Zhang

## Abstract

Tuberculosis (TB) has become a leading cause of death worldwide, which is largely attributed to the difficulties in diagnosis and treatment of TB patients. Exosomes carrying RNA, particularly miRNA, have been indicated their functional and diagnostic potential in diseases, including tuberculosis (TB). In the present study, we performed RNA-seq based analysis on exosomal miRNA profiles for clinical specimens of healthy controls (HC), active tuberculosis (TB) and latent tuberculosis infection (LTBI). We identified many distinct up-regulated and down-regulated differentially expressed miRNA and further screened top 20 in each compared groups which might provide a potential panel for differentiation of HC, LTBI, and TB. We classified all the differentially expressed miRNAs into six expression patterns and identified three persistently up-regulated miRNA (hsa-miR-140-3p, hsa-miR-3184-5p and hsa-miR-423-3p) as potential markers during TB progression. Combined with our previously detected exsomal mRNA, we screened the genes overlapped with predicted mRNA targets of differentially expressed miRNA and analyzed their involvement in Biological Process, indicating a decreased signaling transduction and increased cell death in LTBI and TB. Our results indicate the selective packaging of RNA cargoes into exosomes under different stages of *Mycobacterium tuberculosis* infection and facilitate further study of TB pathogenesis and development.

**IMPORTANCE:** The main reason for failure to eliminate TB is lack of understanding molecular mechanism of TB pathogenesis and difficulties in TB diagnosis and treatment. Exosomes provide a promising research tool because they are released from various cells containing valuable biochemical information related to diseases. We reveale distinct miRNA expression profile of the exosomes, which indicates selective packaging of RNA cargoes into exosomes under different stages of *Mycobacterium tuberculosis* infection. Further, we also provide evidence of related miRNA candidates potentially involving in TB progression and facilitating discovery of TB biomarkers.

## INTRODUCTION

Tuberculosis (TB) is a chronic infectious disease caused by the intracellular bacterial pathogen *Mycobacterium tuberculosis* (*Mtb*). According to WHO report, TB has become a leading cause of death worldwide(1). An estimated 2 billion people (one-third of the world’s population) are latently infected with *Mtb*, yet only 5-10% of the infected individuals will develop active TB during their lifetime, which occurs when the immune response can no longer contain bacilli(2). However, the underlying mechanism for the pathogenesis and progession of TB remains mysterious.

Exosomes are 30-150nm in size and constitutively released from most eukaryotic cell types into the lymphatic system and blood to facilitate cell-cell communication by shuttling various molecules from donor to recipient cells(3). These vesicles carrying cargoes of proteins, lipids and nucleic acids are attributed to cell origin reflecting cellular abnormalities(4), they provide valuable information about disease, including tuberculosis.

Comprehensive proteomic based analysis has characterized the protein cargo of exosomes derived from macrophages infected with *Mtb* or treated with *Mtb* culture filtrate protein as well as from *Mtb*-infected mice(5,6). These exosomes present of host proteins along with mycobacterial proteins have been shown to promote both innate and acquired immune response in vitro and in vivo(5,7–9). Besides, exosomes released from *Mycobacterium avium* infected macrophages contain bacterial pathgenic glycopeptidolipids and cause proinflammatory response(10). Pioneering studies also determined RNA contents in exosomes and the genetic information could exchange between cells(11). These RNAs are highly stable for being protected from enzymic degradation in body fluids(12–14), Which suggests functional and diagnostic potential of exosomal RNAs in tuberculosis. In particular, miRNAs delivered by exosomes have been found to regulate gene expression and cell function both in vivo and in vitro (15–17). Singh and coworkers have characterized signature of host derived miRNAs and mRNA transcripts as well as mycobacterial RNA in exosomes derived from *Mtb* infected macrophages(18). However, RNA-seq based analysis in human clinical specimens is still lacking.

Our previous study have profiled exosomal mRNA signatures in healthy controls (HC), latently infected individuals (LTBI) and patients with active tuberculosis (TB)(19). In the present study, we also applied RNA-seq for small RNA library to further analyze exosomal miRNA profiles of serum samples from HC, LTBI and TB. We revealed distinct miRNA expression profile of the exosomes, indicating the selective packaging of miRNA cargoes into exosomes under different physiological status. We further determined six expression patterns of differentially expressed miRNA and three persistently up-regulated miRNAs (hsa-miR-140-3p, hsa-miR-3184-5p and hsa-miR-423-3p) were screened as potential mediaters during TB progression. Finally we co-analyzed the differentially expressed miRNA with differentially expressed mRNA that we have reported so as to exploring the functional role of exosomal RNA in TB pathogenesis. Our results provided important information of exosomes during Mtb infectious process and encouraged considering exosomal miRNAs as potential biomarkers in TB diagnosis.

## RESULTS

### Baseline characteristics of HC, LTBI and TB

A total of 180 participants were enrolled, including persons with 60 LTBI, patients with 60 TB and 60 healthy control. The age of the LTBI was 40.2±7.9 years old in average, while the mean age was 42.1 ± 8.2 for TB and 38.0 ± 9.1 in HC respectively. Besides, the gender ratio and the number of participants smoking or not was almost the same among groups of HC, LTBI and TB (Table 1), indicating a balanced distribution of baseline characteristics.

**Table 1.** characteristics of the participants

|  | HC | LTBI | TB |
| --- | --- | --- | --- |
| n | 60 | 60 | 60 |
| Male/Female | 24/36 | 27/33 | 22/37 |
| Mean age ± SD(y) | 38.0±9.1 | 40.2±7.9 | 42.1±8.2 |
| Age range(y) | 20-55 | 23-56 | 19-65 |
| Smokers/Nonsmokers | 15/45 | 18/42 | 19/41 |

Note: HC: healthy control, LTBI: latent tuberculosis infection, TB: tuberculosis. Serum exosomes were obtained from pools composed of 60 participants for the HC, LTBI and TB samples, respectively.

### Expression profiles of exosomal miRNA from serum of TB, LTBI and HC

Small RNA reads were obtained using high-throughput sequencing strategy. By mapping to the human reference genome (hg38), we obtained the small RNA expression profiles of exosomes derived from the three groups (HC, LTBI and TB). A total of 9584942 (HC), 778169 (LTBI), and 10581000 (TB) reads were identified, of which 8681012 (90.57%), 688360 (88.46%), and 3370891 (31.86%) small RNAs were identified in HC, LTBI, and TB respectively (Supplementary Table S1). Via aligning sequences with the miDeep2 database, we identified a total of 435 miRNAs, among which 364 sequences were consistent with mature miRNA records in miRBase, while 71 sequences that not paired to any records were predicted as novel miRNAs (Supplementary Table S2). Taking all the aligned miRNAs into consideration, we found that the expression value of miRNAs presented a decreased trend in LTBI compared to HC, and TB has the lowest expression level among the three goups (Figure 1a), which was in accordance with Singh PP’s discovery that incorporation of miRNA into exosomes was inhibited in Mtb infected mouse macrophages but not so in non infected control cells(18).

**Figure 1.**
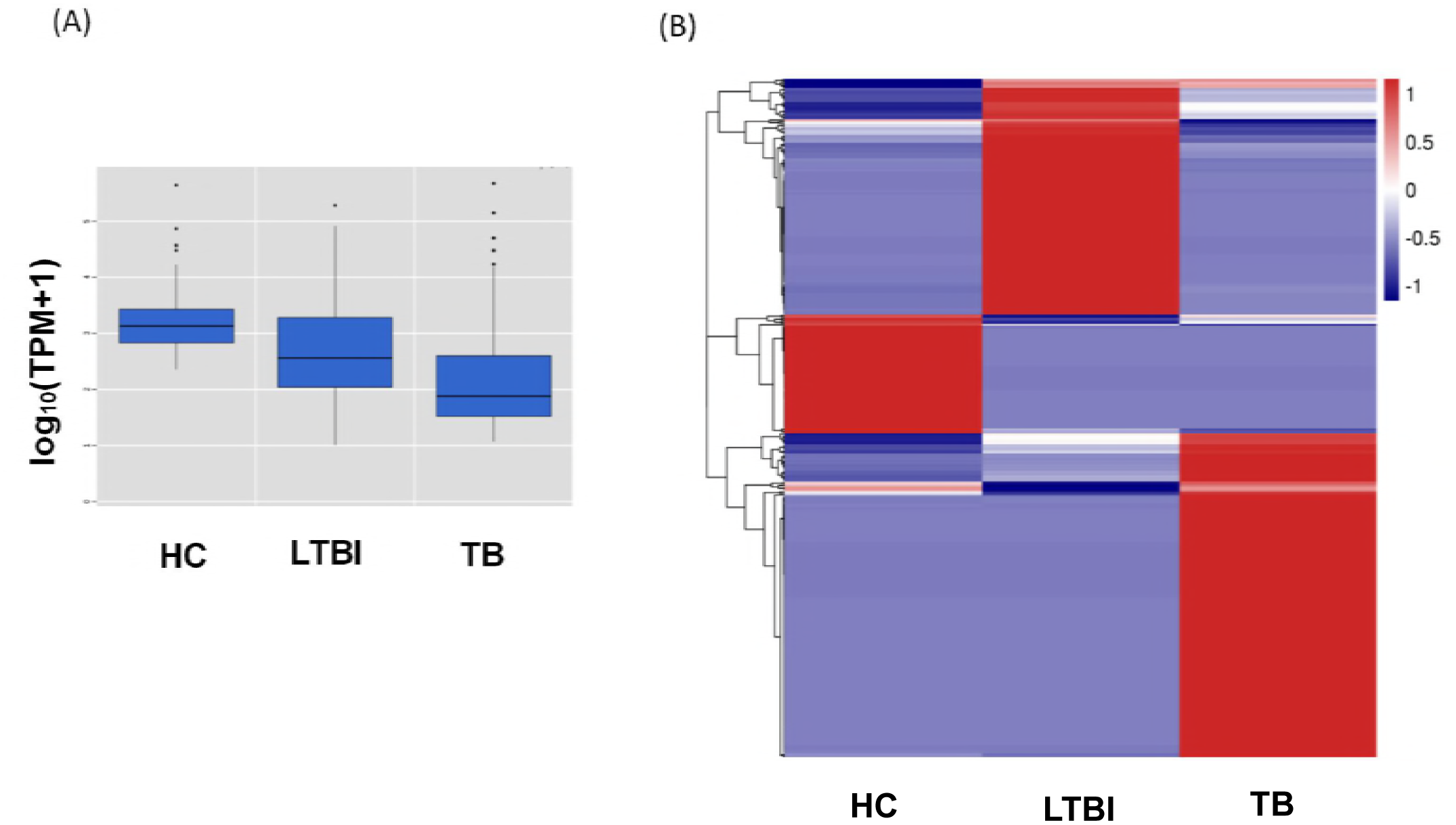
Expression profile of miRNA in serum exsomes derived from HC, LTBI and TB. **(A)** The box plot used to show the TPM values of miRNA in each sample. **(B)** Heatmap of the expression of different miRNAs (red, high expression; blue, low expression)

### Differential miRNA expresstion profiles among the serum exosomes in the HC, LTBI and TB

To identify the significant differenecs of miRNAs between either two groups, we profiled the differential miRNAs by pairwise comparison (fold change ≥2 and q value≤0.05) in a heatmap (Figure 1b). The results revealed the distinct miRNAs expression profiles of the exosomes from HC, LTBI, and ATB: 32 miRNAs were highly expressed in HC samples compared with LTBI and ATB samples (the left column of Figure 1b), 40 miRNAs (the middle column) were highly expressed in LTBI samples, while 54 miRNAs (the right column) were highly expressed in TB samples compared with the other two groups. Notably, 153 miRNAs were uniquely expressed in TB while 43 miRNAs were specifically identified in LTBI (Supplementary Table S3), which is helpful to facilitate the development of potential targets for the diagnosis of tuberculosis.

Further, fold changes of the differentially expressed miRNAs were analyzed among the three groups. In comparison with the HC samples, we identified 53 up-regulated miRNAs and 66 down-regulated miRNAs in LTBI, while up-regulated miRNAs increased to 63 in TB and down-regulated miRNAs decreased to 36. When TB and LTBI were compared, 114 miRNAs were identified up-regulated and 67 miRNAs were down-regulated (Figure 2a and Supplementary Table S4). We showed the top-10 up-regulated and top-10 down-regulated miRNAs in the above compared groups (Figure 2c). To further investigate the detailed differences in the differentially expressed miRNAs between LTBI and TB, we overlapped these miRNAs using a Venn diagram (Figure 2b). They exhibited relatively distinct expression panels: 43 and 54 miRNAs were only up-regulated and expressed in LTBI and ATB, respectively; 34 and 3 genes were uniquely down-regulated and expressed in LTBI and ATB, respectively; while only 1 miRNA was up-regulated and expressed in LTBI but down-regulated and expressed in ATB. In addition, there was a portion of differentially expressed miRNAs (fold change >2 and q-value <0.05) in ATB and LTBI that shared similar expression panels (9: up-regulation; 32: down-regulation).

**Figure 2.**
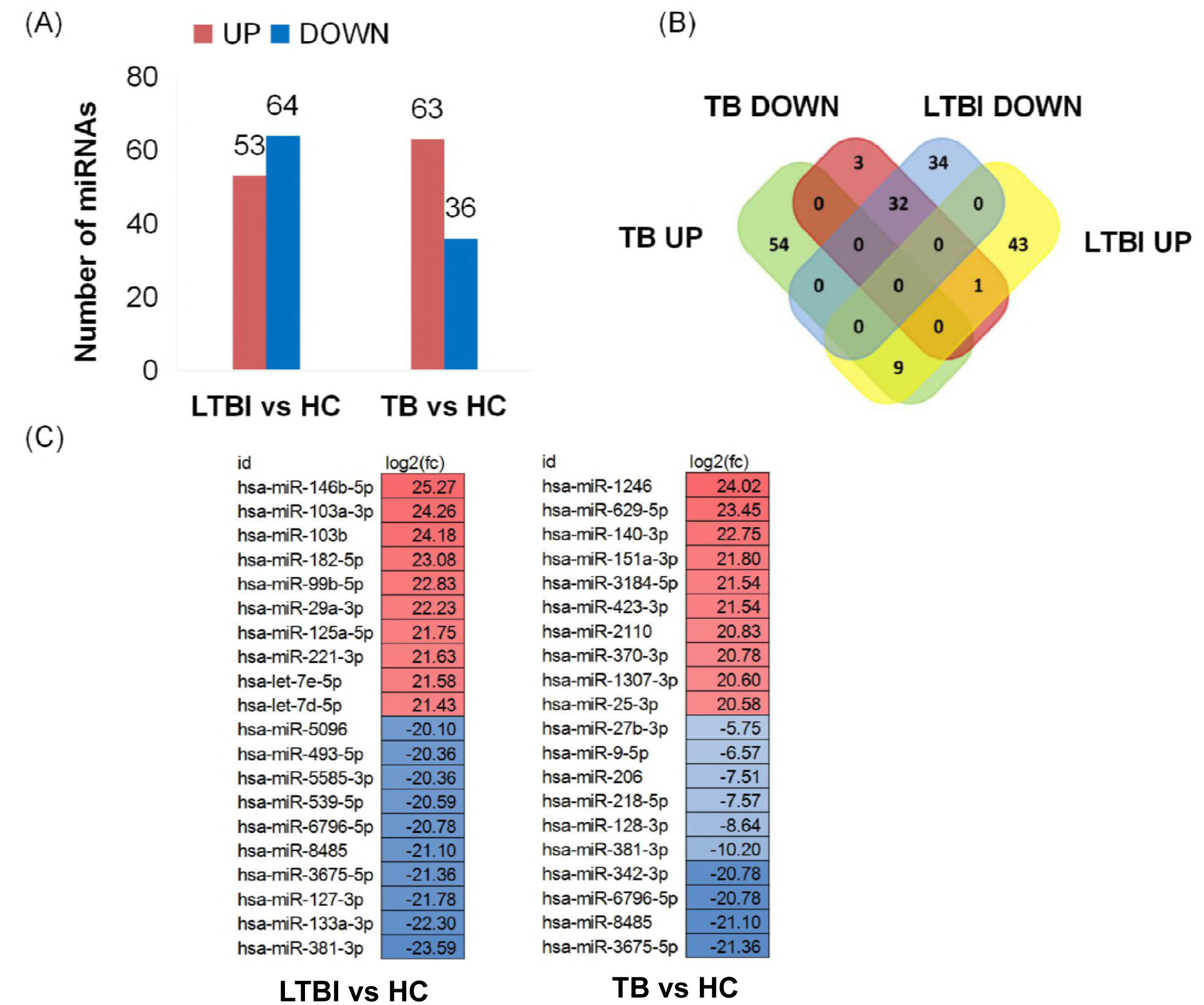
Differential expression of exosomal miRNA in compared groups of LTBI vs HC and TB vs HC. **(A)** Bar plot of the numbers of the differentially expressed miRNAs in LTBI and TB compared with HC. Red bars represent the number of up-regulated genes in each sample, while the blue bars represent the number of down-regulated genes in each sample. **(B)** Venn diagram of differentially expressed miRNA showing the relatively distinct expression panels between the TB and LTBI. **(C)** Top 10 up-regulated miRNA and down-regulated miRNA in three samples, respectively.

### Expression patterns of serum exosomes in HC, LTBI, and TB individuals

We classified all the differentially expressed miRNAs into six expression patterns according to the gene expression trends among the three samples, including MLH, MHL, LMH, LHM, HML and HLM (L: low expression level, M: mediaum expression level, H: high expression level) (Figure 3). Hight expressed miRNAs in LTBI and TB which were marked with a red box in Figure 3 provided potential targets of clinical diagnosis for these two groups of individuals. We next examined the gradually up-regulated or down-regulated miRNAs which might play important roles in TB pathogenesis. There were three miRNAs (hsa-miR-140-3p, hsa-miR-3184-5p and hsa-miR-423-3p) significantly up-regulated in LMH pattern but no miRNA was obviously down regulated in HML pattern from HC to LTBI and then to TB.

**Figure 3.**
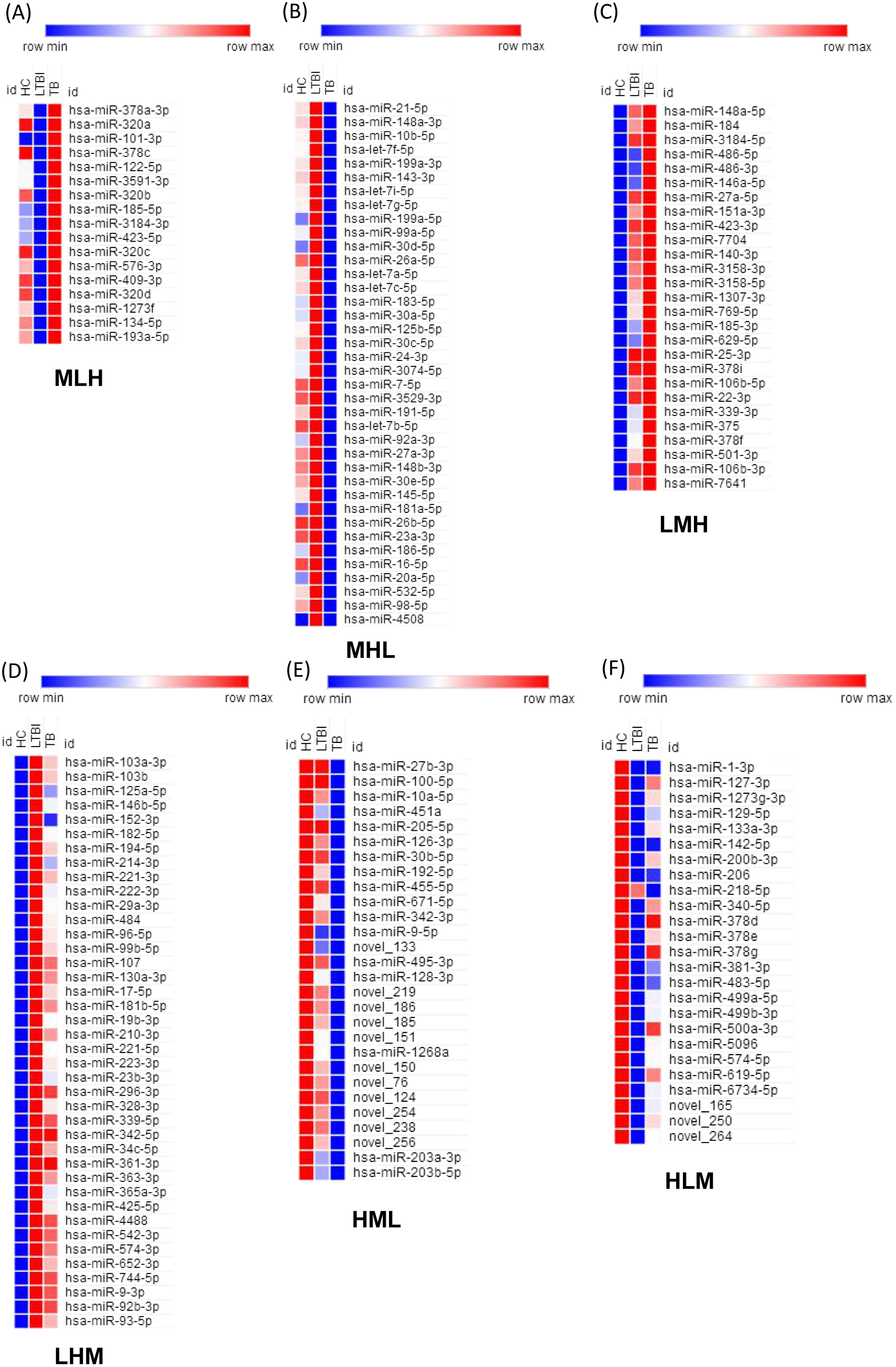
Expression patterns of exosomal miRNA among groups of HC, LTBI and TB. Heatmaps used to show six patterns of miRNA expression based on TPM values, including **(A)** MLH, **(B)** MHL, **(C)** LMH, **(D)** LHM, **(E)** HML, **(F)** HLM. (L: low expression level; M: medium expression level; H: high expression lesvel).

Analysis of the three miRNAs with miRPath (v3.0) showed that combinations of them overlapped in functional catorgies such as fibroblast growth factor receptor signaling pathway, neurotrophin TRK receptor signaling pathway, Fc-epsilon receptor signaling pathway, cellular protein modification, small molecule metabolic process, iron binding, enzyme binding, blood coagulation and organelle (Figure 4), and they also overlapped in enriched pathways including ECM-receptor interaction, phototransduction, axon guidance, Ras signaling pathway, Rap1 signaling pathway, bacterial invasion of epithelial cells, huntington’s disease (Table 2).

**Figure 4.**
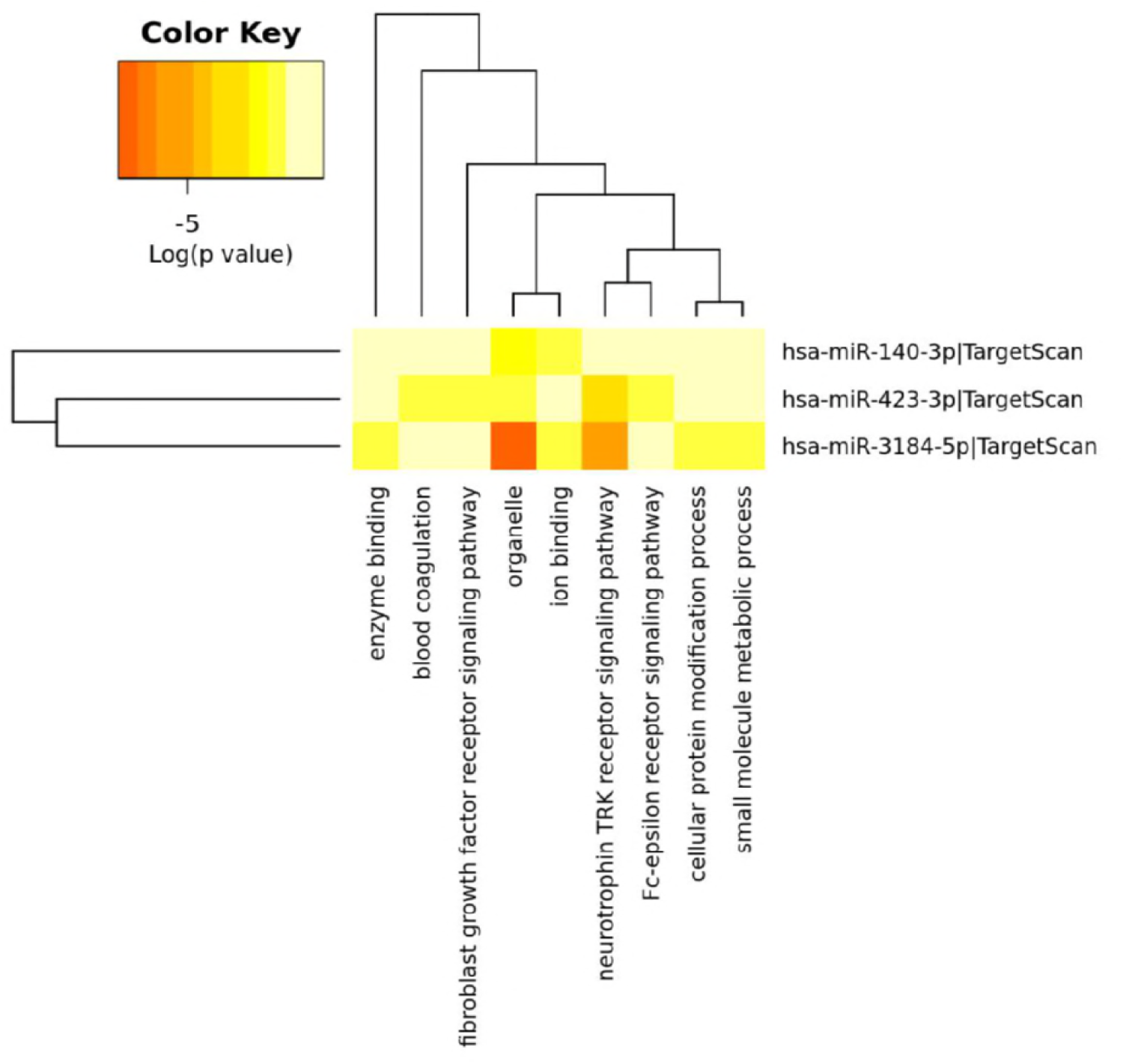
Heatmap of significantly up-regulated miRNA between compared groups of LTBI vs HC and TB vs LTBI, and their predicted GO items. Heat map showed the predicting GO catagories of three persistantly up-regulated miRNA in TB development. The heatmap was generated using mirPath v.3 (Diana Tools) as described.

**Table 2.** Predicted KEGG pathways of genes targeted by mihsa-miR-140-3p, hsa-miR-3184-5p and hsa-miR-423-3p

| KEGG pathway | p-value | #genes |
| --- | --- | --- |
| ECM-receptor interaction | 1.56E-07 | 7 |
| Phototransduction | 0.000555956 | 6 |
| Axon guidance | 0.004731024 | 11 |
| Ras signaling pathway | 0.005979265 | 22 |
| Rap1 signaling pathway | 0.022264876 | 20 |
| Bacterial invasion of epithelial cells | 0.045859124 | 9 |
| Huntington's disease | 0.048087402 | 8 |

Note:
#: number

### miRNA targets converge on functional annotation with published exosomal transcriptome in HC, LTBI and TB

The regulatory genes of distinctive miRNA in LTBI vs HC and TB vs HC were predicted using TargetScan and MicroT-CDs database. Overlapped genes and validated mRNA targets from miTarbase were unioned as input data into DAVID 6.8 (cutoff was set as p-value≤0.05) for further functional annotation. The Top 10 enriched GO catagories for each aspect were shown in Figure 5, and some GO items, for example, cellular nitrogen compound metabolic process and iron binding that were functionally relevent for Mtb pathogenesis(20, 21). On the other hand, the KEGG pathway analysis (Top 20 were shown in Figure 5) indicated those miRNA targeted genes were invovled in signaling pathways associated with immune response to Mtb infections, such as MAPK signaling pathway, TGF-beta signaling pathway, Wnt signaling pathway, Endocytosis, Hippo signaling pathway and so on(22–25).

**Figure 5.**
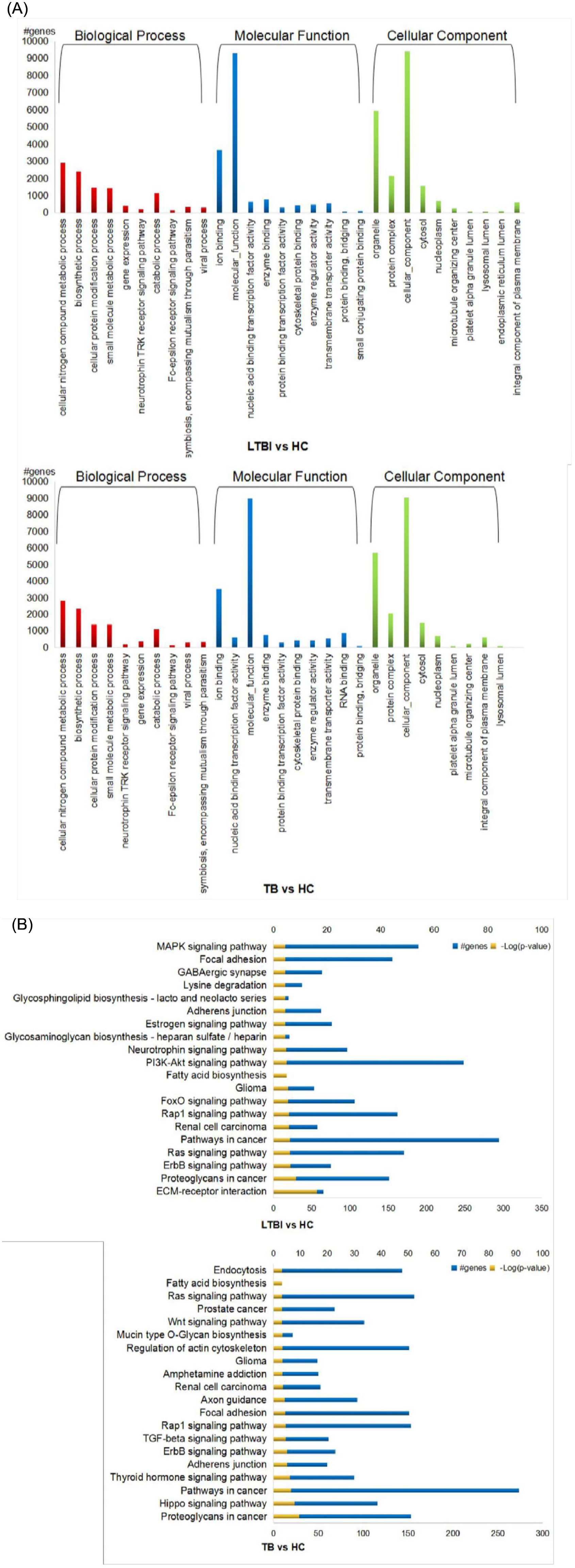
GO annotation KEGG pathway of differentially expressed miRNA targeted genes in compared groups of LTBI vs HC and TB vs HC. **(A)** The top 10 GO items for biological process, molecular function, and cellular component for distinctive miRNAs in LTBI vs HC and TB vs HC. **(B)** The KEGG pathway enrichment of distinctive miRNAs in LTBI vs HC and TB vs HC.

Our previous published data comparing the transcriptome of exosomes derived from HC, LTBI and TB were engated into analysis together with the differentially expressed exosomal miRNA targets in this study. In comparison with HC, we have identified 769 up-regulated mRNAs and 643 down-regulated mRNAs in the LTBI individuals, while it was 999 up-regulated and 1582 down-regulated mRNAs in TB individuals respectively (Supplementary Table S5). If mRNA target repression by miRNAs is assumed, up-regulated transcripts would be the result of decreased miRNA and down-regulated transcripts the result of increased miRNA. We further found that 167 up-regulated transcripts overlapped with the mRNA targets of our down-regulated miRNAs and 164 down-regulated transcripts overlapped with targets of up-regulated miRNAs in LTBI compared with HC. While the number was 220 and 582 in TB compared with HC, respectively (Figure 6 and Supplementary Table S5). Then we used the overlapped mRNA in each compared groups as input into FunRich_V3 (cutoff was set as p-value≤0.05). The results showed that down-regulated mRNAs in LTBI were mainly enriched in Biologic Process (BP) of cell growth and/or maitenance, regulation of gene expression/epigenetic, cell motility, regulation of endocytosis and synaptic transmission. Meanwhile, the diminished BP items were quite different in TB samples, which included regulation of nucleobase,nucleoside,nucleotide and nuleic acid metabolism, regulation of cell cycle, electron transport, cell-cell adhesion, anti-apoptosis, cell migration as well as regulation of gene exression. When examining up-regulated BP catorgries, there were nuclear organization and biogenesis, gene silencing hormone metabolism, cell differentiation, lipid metabolism, in addtion to unknown biological process in LTBI, while the robust BP items in TB were transport, cell organization and biogenesis, homeostasis, regulation of blood pressure and skeletal development.

**Figure 6.**
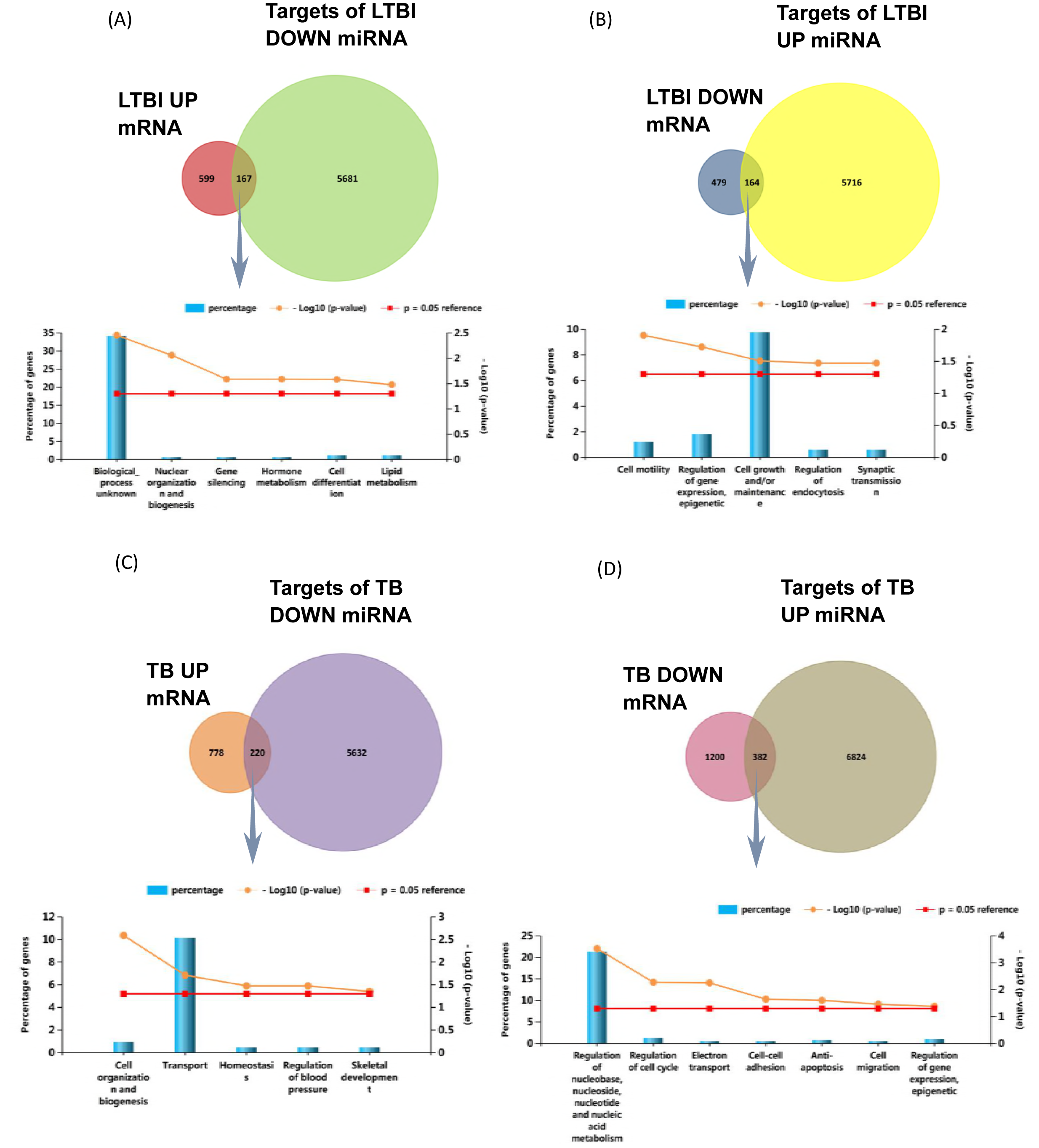
Overlapped genes of predicted mRNA with experimentally detected exosomal mRNA and their involvement in Biological Process in compared groups of LTBI vs HC and TB vs HC. Venn diagram showed the overlapped genes of **(A)** up-regulated mRNA with predicted mRNA targets of down-regulated miRNA in LTBI. **(B)** down-regulated mRNA with predicted mRNA targets of up-regulated miRNA in LTBI. **(C)** up-regulated mRNA with predicted mRNA targets of down-regulated miRNA in TB. **(D)** down-regulated mRNA with predicted mRNA targets of up-regulated miRNA in TB. while the bar plot showed those overlapped genes’ enriched items of GO items for biological process, respectively.

### Mtb RNAs were detected in LTBI and TB samples

Previous study have indicated that *Mtb* components including peptides and RNAs were found in exosmes derived from *Mtb* infected host cells (5,6,18). We performed Blast analysis against *Mtb* reference genome (NC00962) using the sequenced small RNA for the three groups. As shown in the Supplementary Table S6, 286 RNAs and 11309 RNAs were identified in LTBI and TB, respectively. Surprisingly, there were 227 RNAs mapped to the reference genome in HC, Which was in consistent with our previous discovery and Gutkin’s finding that a few *Mtb* RNA and peptides were observed in HC individuals(19, 26).

## DISCUSSION

To date, TB is still a serious threat to human health. The main reason for failure to eliminate TB is lack of understanding the molecular mechanism of TB pathogenesis and progression. In recent years, exosomes have been shown to participate in signal transduction (e.g. immune regulation)(27), biomolecules transportation (e.g. nucleic acids, proteins, lipids)(28), and cellular "trash bags" for elimination of excess intracellular substances(29). Studies on exosomal RNAs in immunology, cancer biology and neurobiology have indicated the potential use of them as molecular targets against several diseases(30–33). However, the role of exosomes in transporting genetic material, specifically miRNA, in tuberculosis with clinical human species was still being defined.

In the present study, we identified many distinct up-regulated and down-regulated differentially expressed miRNA and screened the top 10 miRNA in each panel from the three groups. Previous studies have indicated differentially expressed miRNAs in clinical serum/plasma as potential biomarkers for TB diagnosis. Since majority of miRNAs in human serum or saliva are concentrated in exosomes allowing to protected from enzymatic degradation(34), we examined our analytic results by comparing to theirs’. Qi and colleagues reported that miR-576-3p could differentiate TB patients from HC with moderate sensitivity and specificity(35). Consistently, we also demonstrated miR-576-3p were obviously up-regulated in TB patients when compared to HC. However, we showed down-regulation of miR-483-5p and up-regulation of miR-486-5p in TB patients in our study, which was reverse in other published studies (36,37). Besides, couples of differentially expressed miRNAs, such as miR-93(38), miR-29a(38), miR-378(36), miR-22(36), miR-196b(37) and miR-155(39), were reported to serve as potential diagnostic markers for TB, some miRNAs in our study belonging to the same miRNA family, including miR-93-5p, miR-29a-3p, miR-378d, miR-378f, miR-378i, miR-22-3p, miR-196b, miR-155-5p, were also detected their expression in the same regulatory pattern in TB patients. Additionally, miR-let-7e-5p was highly up-regulated in LTBI compared to HC, but it became sharply down-regulated in TB compared to LTBI. MiR-let-7 family is well known for their role in apoptosis of cancer cells and recently was reported their novel function in regulation of anti-*Mtb* immune response(38,40). Together with a recent report that exosome-enclosed miRNAs in exhaled breath has been indicated to hold potential for biomarker discovery in patients with pulmonary diseases, including tuberculosis(41), these distinct miRNA shed lights on discovery of novel biomarkers for LTBI and TB diagnosis.

Besides, we classified all the differentially expressed miRNAs into six expression patterns. In order to discover the mechanism of RNA packaging into exosomes during disease progression, we focused on the LMH pattern and HML pattern, which presented grandually up-regulated and down-regulated miRNA respectively under status as Mtb infection progresses. We found three persistently up-regulated miRNA including hsa-miR-140-3p, hsa-miR-3184-5p and hsa-miR-423-3p were significantly distinct (q-value≤0.05 and |log2(foldchange)|>1) in LTBI vs HC and TB vs LTBI, but no obvious down-regulated miRNA were determined, indicating the inhibitory role of exosomes in TB development. However, the specific role and the regulatory mechanism of three miRNA candidates in disease progression need further identificaiton.

By co-analyze miRNA in this study with our previous mRNA data based on the assumption that the expression level of miRNA and its targeted mRNA were mostly reverse, we screened the genes overlapped with predicted mRNA targets of differentially expressed miRNA and analysed their involvement in Biological Process. Our results indicated a decreased signaling transduction and increased cell death in Mtb infected individuals, especially in TB patients.

Interestingly, we also detected mycobacterial RNA fragments in exosomal samples derived from LTBI and TB, and the amount of *Mtb* small RNA was much more in TB (11309, 0.11%) than that in LTBI (286, 0.04%), which was opposite to our published result that Mtb transcripts mostly existed in exosomes of LTBI rather than that of TB. Taken together, it was suggested that exosomes derived from LTBI which contained more Mtb transcripts might play a regulatory role in Mtb survival in recipient cells while exosomes from TB pattients that contained more sRNA might act as a role in cellular apoptosis, since mycobacterial sRNA has been reported to induce apoptosis in human macrophages(42). How mycobacterial RNAs are incorporated into the exsomes is presently unclear. Previous study has shown that mycobacterial DNA get access to cyosolic receptors by perforating the phagosome membrane mediated by the ESX-1 secretion system(43). Since we only generated small RNA library for sequencing, the length of *Mtb* RNA fragments are unclear and need further validation for further use of this bacterial-derived RNA in TB diagnostics.

In conclusion, our study demonstrated that miRNA expression profiles were significantly different in exosomes isolated from serum of HC, LTBI and TB. Couples of differentially expressed miRNAs in TB or LTBI had the potential to serve as biomarkers for TB diagnosis. Three miRNA were proposed to be mediators in TB development. *Mtb* sRNA were incorporated into exosomes from TB. However, confirmation of these potential RNAs in a clinically relevant sample set will offer a source for novel TB biomarkers and advance our knowledge about TB pathogenesis or progression.

## MATERIALS AND METHODS

### Study participants

The study was conducted in accordance with the principals of the Declaration of Helsinki and approved by the Ethical committees of Beijing Chest Hospital, Capital Medical University. All participants were at least 18 years old, HIV-negative and written, informed consents were obtained from them. TB patients were classified based on having clinical presentation compatible with TB infection, a positive *Mtb* culture and a positive smear. Patients were excluded if they had previous TB history or had received anti-TB therapy before enrollment. Latently infected subjects defined as culture negative, having positive tuberculin skin test (TST) and interferon-gamma release assay (IGRA) using T-SPOT.TB (Oxford Inmmunotec, Abingdon, UK), normal chest computed tomography (CT), no clinical symptoms or evidence of active TB and other non-tuberculosis respiratory infections. The TST/IGRA two-step strategy is used because the confirmatory IGRA is able to highly reduce the false positivity due to BCG vaccination or nontuberculous mycobacteria infection (NTM) in initial TST. Healthy *Mtb* uninfected controls were enrolled with negative TST and T-SPOT.TB tests, normal chest CT and no clinical evidence of any diseases. The participants demographic information is shown in table 1.

### Sample preparation

A total of 180 serum samples were collected. The serum samples were grouped according to the clinical cohort as active tuberculosis (TB), Latent tuberculosis infection (LTBI) and healthy control (HC). Serum was obtained from each participant and then pooled based on the group (pooled n=60 for each pooled sample).

### Exosome isolation and characterization

Exosome isolation from pooled serum samples was conducted as previously described. Briefly, the cell debris were removed by differential centrifugation at 1000 g for 10 min at 4°C, and 16,500 g for 30 min at 4 °C, followed by ultrafiltration (through a 0.22 μm filter; Millipore, Billicera, MA, USA). Then, the exosome pellet was obtained by ultracentrifugation at 120,000 g for 2 hrs) and washed with PBS.The pellets was resuspended in water and dropped on 200 mesh copper grids, the fluid was then absorbed from the edges of the copper mesh with filter paper. The copper mesh was stained with Phosphotungstic acid for 10 min, washed 3 times with water, and then dried for observation by transmission electron microscopy (TEM, FEI, Netherlands). By using the nanoparticle tracking analysis (NTA, Malvern, UK) according to the manufacturer’s protocol, an absolute size distribution of the exosmes was obtained (Supplemental Figure 1).

### Exosmal RNA extraction

Isolated exosomes were immediately used for total RNA extraction using RNAiso-Plus (TaKaRa, Dalian, China) according to the manufacturer’s instructions. RNA concentration was measured using Qubit® RNA Assay Kit in Qubit® 2.0 Flurometer (Life Technologies, CA, USA). RNA integrity was assessed using the RNA Nano 6000 Assay Kit of the Agilent Bioanalyzer 2100 system (Agilent Technologies, CA, USA).

### Sequencing and data processing

Sequencing libraries were generated using NEBNext® Ultra^™^ Directional RNA Library Prep Kit for Illumina® (NEB, MA, USA) following manufacturer’s recommendations and index codes were added to attribute sequences to each sample. The library quality was assessed on the Agilent Bioanalyzer 2100 system (Agilent Technologies, CA, USA). The RNA libraries were sequenced on the Illumina Hiseq 2500/2000 platform and 50 bp single reads were generated.

The clean small RNA reads were mapped to human reference genome (hg38) by Bowtie(44) without mismatch to analyze their expression and distribution on the reference. To remove tags originating from protein-coding genes, repeat sequences, rRNA, tRNA, snRNA, and snoRNA, small RNA tags were mapped to miRNA database (miRBase, http://www.mirbase.org/). miRNA expression levels were estimated by TPM (transcript per million) through the following criteria(45): Normalization formula: Normalized expression = mapped read count/Total reads^*^1000000. For the samples with biological replicates: Differential expression analysis of two conditions/groups was performed using the DEGseq (2010) R package. P-value was adjusted using q-value(46). q-value≤0.05 and |log2(foldchange)|>1 was set as the threshold for significantly differential expression by default.

### miRNA target gene prediction and functional annotation

Predicting the target gene of miRNA was performed by microT-CDs database (http://diana.imis.athena-innovation.gr/DianaTools/) with a miTG score ⩾ 0.8 set, and TargetScan (http://www.targetscan.org/vert_72/), the overlapped mRNAs were then processed full outer join with the experimentally validated target genes by miRTarBase (http://mirtarbase.mbc.nctu.edu.tw/php/index.php) with methods of immunoprecipitation, western blot, qPCR, reporter gene assay, microarrays, or others were labeled, respectively. Functional annotation of targeted genes was conducted using GO analysis and KEGG pathway enrichment via DAVID 6.8 online (https://david.ncifcrf.gov/). The heatmap of GO terms and clustering of predicted KEGG pathways of persistantly up-regulated three miRNAs were performed according to DIANA mirPath v.3(47). The functional distribution of mRNA candidates according to Biological Process were based on information provided by the online resource FUNRICH system(48).

## ACKNOWLEDGEMENTS

This work was supported by Beijing Natural Science Foundation (5174035), National Natural Science Foundation of China (31700668), National Science and Technology Major Project (2017ZX10201301-004 and 2015ZX10004801-003), Collaborative Innovation Center of Infectious Diseases (PXM2016_014226_000052), Beijing Municipal Administration of Hospitals’ Ascent Plan (DFL20181601), and Tongzhou District Science and Technology Committee (KJ2017CX076).

## ETHICAL APPROVAL

This study was carried out in accordance with the recommendations of the Helsinki Declaration and its later amendments or comparable ethical standards, the Ethics Committee of the Beijing Chest Hospital, Capital Medical University. This article does not contain any studies with animals performed by any of the authors. The protocol was approved by the Ethics Committee of the Beijing Chest Hospital, Capital Medical University.

